# A versatile automated high-throughput drug screening platform for zebrafish embryos

**DOI:** 10.1101/2020.12.16.423108

**Authors:** Alexandra Lubin, Jason Otterstrom, Yvette Hoade, Ivana Bjedov, Eleanor Stead, Matthew Whelan, Gaia Gestri, Yael Paran, Elspeth Payne

## Abstract

Zebrafish provide a unique opportunity for drug screening in living animals, with the fast developing, transparent embryos allowing for relatively high throughput, microscopy-based screens. However, the limited availability of rapid, flexible imaging and analysis platforms has limited the use of zebrafish in drug screens. We have developed a easy-to-use, customisable automated screening procedure suitable for high-throughput phenotype-based screens of live zebrafish. We utilised the WiScan^®^ *Hermes* High Content Imaging System to rapidly acquire brightfield and fluorescent images of embryos, and the WiSoft^®^ *Athena Zebrafish Application* for analysis, which harnesses an Artificial Intelligence-driven algorithm to automatically detect fish in brightfield images, identify anatomical structures, partition the animal into regions, and exclusively select the desired side-oriented fish. Our initial validation combined structural analysis with fluorescence images to enumerate GFP-tagged haematopoietic stem and progenitor cells in the tails of embryos, which correlated with manual counts. We further validated this system to assess the effects of genetic mutations and x-ray irradiation in high content using a wide range of assays. Further, we performed simultaneous analysis of multiple cell types using dual fluorophores in high throughput. In summary, we demonstrate a broadly applicable and rapidly customisable platform for high content screening in zebrafish.

## Introduction

Zebrafish provide an excellent model for human disease and offer a unique opportunity for *in vivo* small-molecule phenotypic drug screening. Each breeding pair can lay hundreds of embryos, which combined with the rapid development and transparent nature of the embryos makes them amenable to microscopy-based screens usually otherwise restricted to cell culture. Unlike most *in vivo* screening platforms, thousands of animals can be imaged within days, allowing for a relatively high-throughput screen, with the advantages of screening in intact living animals.

The utility of such *in vivo* screens has been demonstrated by the rapid repurposing of identified compounds into clinical trials. This is exemplified by dmPGE2, which entered clinical trials as a therapy for patients undergoing umbilical blood cord transplantation, having been found to enhance haematopoietic stem cells in a zebrafish screen using *in situ* hybridization (North et al., 2007, Hagedorn et al., 2014). Additionally, ORC-13661, identified in zebrafish screens of hair cells in zebrafish embryos (Owens et al., 2008, Chowdhury et al., 2018, Kitcher et al., 2019) is currently in clinical trials as an agent to prevent hearing loss from aminoglycoside antibiotic-induced hair loss.

Despite their advantages, the limited availability of image acquisition and, especially, analysis platforms supporting zebrafish in a fast, flexible format has limited their widespread uptake in drug screens. Screens are typically slow, often relying on manual or bespoke imaging solutions and manual analysis. For example, North *et al.* utilised a manual qualitative scoring after *in situ* hybridization using two probes (North et al., 2007), to identify dmPGE2. The discovery of ORC-13661 as a modifier of hair cells also relied on manual inspection of fluorescent neuromasts for 10,960 compounds after treatment with a dye, with manual counting required to quantify changes (Owens et al., 2008). This hair cell assay is rapid and simple and has been used in a number of additional screens (Chiu et al., 2008, Vlasits et al., 2012, Esterberg et al., 2013, Pei et al., 2018), however all still rely on manual counting or scoring of individual fish, creating a significant bottleneck in analysis throughput.

The optimal zebrafish high-throughput screening platform would permit automation of both image acquisition and analysis across a range of multiplexed assays and phenotypes with minimal human intervention. One of the primary challenges in this process is getting the embryo in the desired orientation for imaging without manual manipulation of the embryos. One approach has been to use small glass capillaries for imaging embryos, as in the VAST Bioimager which allows for automated imaging of zebrafish embryos in a chosen orientation (Pulak, 2016, Pardo-Martin et al., 2010, Pardo-Martin et al., 2013, Chang et al., 2012). Zebrafish embryos can also be imaged in a semi-automated format by standard microscopes in 96-well plates (Romano and Gorelick, 2014), although there is limited control over the orientation of the fish without manual manipulation or inspection of the images.

Existing automated image analysis solutions are much more limited, with most platforms developed as bespoke solutions. One popular technique is to design algorithms to automatically analyse fluorescent images of transgenic zebrafish, or fish fluorescently labelled with a dye. A number of such screens have adapted the ImageXpress High-Content Screening System by Molecular Devices for automated image acquisition. This has included assays to quantify the number of angiogenic blood vessels (Tran et al., 2007), to analyse axon length (Kanungo et al., 2011) and to measure tumour size (Zhang et al., 2014). However, the automated analysis for these has often required either the development of bespoke algorithms, or the custom adaption of more general software. There are a number of opensource approaches that have been designed or adapted for analysis of zebrafish embryos. QuantiFish is an opensource application developed for analysing fluorescent foci in zebrafish, which has been used to analyse bacterial infection in embryos (Stirling et al., 2020). CellProfiler is a more general image analysis platform (Kamentsky et al., 2011), which has a number of published pipelines including for the analysis of zebrafish embryos to quantify haemoglobin (Metelo et al., 2015). However, to adapt these platforms for new applications requires a lot of development by the user to set up the analysis, which can be challenging without expertise, limiting widespread usage. Artificial Intelligence (AI) based approaches can provide the opportunity for automated image analysis with broader applicability. Using Definiens Cognition Network Technology (CNT), Vogt et al. designed and trained an algorithm to detect and segment transgenic fluorescent embryos arrayed in 96-well plates and quantify blood vessel development (Vogt et al., 2009). They were then able to adapt this method to a different transgenic fish line and phenotype, which was used in a chemical screen for FGF signalling (Vogt et al., 2010, Saydmohammed et al., 2011).

We sought a screening platform to conduct small-molecule and genetic screens looking at haematopoietic stem and progenitor cells (HSPCs) in zebrafish embryos. We set out to develop an automated screen, with a long-term goal of screening for targeted therapeutics for myeloid malignancies. The Tg(itga2b:GFP) transgenic line, which labels thrombocytes and HSPCs with green fluorescence (Lin et al., 2005), provides a readout of the number and location of HSPCs. A semi-automated screen for HSPCs has previously been developed (Arulmozhivarman et al., 2016), but this screen required a custom analysis platform and user-input to define the region of interest.

We have developed an easy-to-use, yet broadly applicable zebrafish embryo screening platform automating both image acquisition and quantitative analysis. We initially performed phenotypic validation for our screen of interest defining HSPCs using genetic and radiation screens of known phenotype. We then extended our assays to combine with multiple fluorescent outputs using mCherry-tagged myeloid cells, and further expanded our screen readout to include the apoptotic acridine orange for use in toxicity screens. To highlight the breadth and simplicity of the platform we also utilised a hair cell marker that has previously been used in a number of manual screens, and a brightfield morphological screen of eye size. Our customisable screen can easily and rapidly be applied to study of other anatomical sites and phenotypes for generalized drug screening in zebrafish, with automated image acquisition and analysis of thousands of fish in a single day.

## Results

### Automatic detection of zebrafish embryos and internal anatomy in multiplexed fluorescence imaging with the co-developed WiSoft^®^ Athena image analysis platform

Effective high-content screening (HCS) necessitates simple and fast image acquisition to allow for high throughput. We utilised the WiScan^®^ *Hermes* High Content Imaging System (IDEA Bio-Medical) to rapidly acquire both fluorescent and brightfield images of live zebrafish embryos at 3dpf. The workflow is depicted in Figure 1A. Live phenylthiourea-treated embryos were anaesthetised and loaded into a 96-well zebrafish alignment plate (Hashimoto) with one embryo per well, using a manual pipette. The plate was briefly centrifuged, before imaging with the *Hermes* (Fig. 1A). Embryos were imaged using a 4X objective and Z-stack acquisition of five slices, spanning 0.2 mm, with four overlapping images along each well to permit accurate image stitching and full-fish visualisation (Fig. S1). Image acquisition was carried out in brightfield and fluorescent channels, taking around 15 minutes to obtain the 3,840 raw images per full plate, allowing for the relatively high throughput. Image pre-processing steps are carried out automatically by the accompanying batch image processing software package (IDEA Bio-Medical, see Methods) in around 20 minutes. We performed best z-slice selection (most in-focus z-plane) for the brightfield channel and maximum z-axis intensity projection for fluorescence channels prior to image stitching for full-fish analysis (Fig. S1), with images processed in batches by accompanying software. Fluorescence max-intensity was chosen over best-slice selection to capture all fluorescently labelled cells throughout the fish volume. In this fashion we were able to visualise individual fluorescently labelled cells without the need for additional processing or image deconvolution while also having both the tail and head of the fish in proper focus for analysis of the brightfield channel.

**Figure 1.**
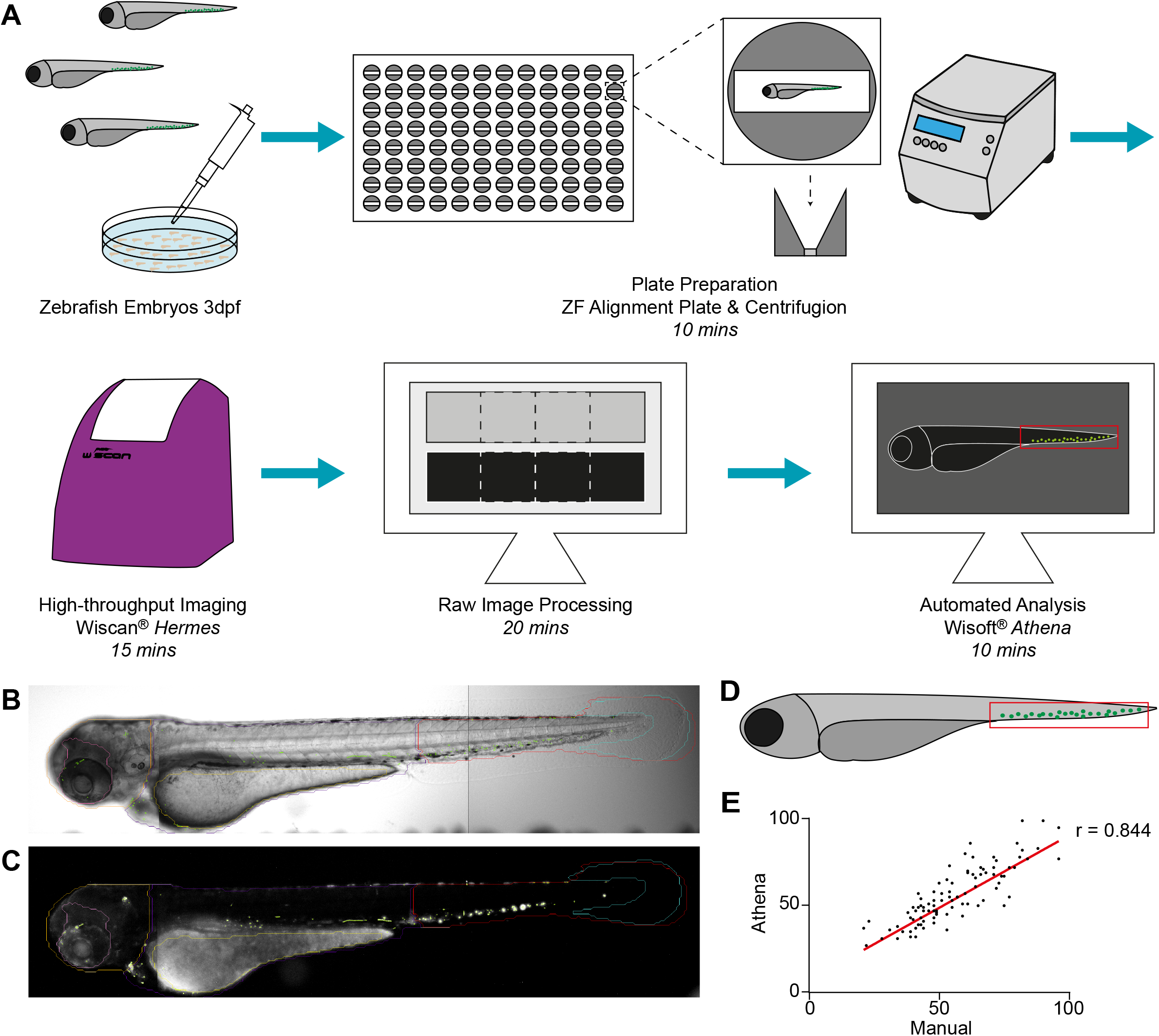
(A) Schematic of screening platform workflow including plate preparation, image acquisition and image analysis of zebrafish embryos with approximate timings. (B) Brightfield image and (C) fluorescent image of a Tg(itga2b:GFP) zebrafish at 3dpf acquired from the *Hermes* showing some of the regions identified by *Athena*: tail (red), trunk (purple), head (orange), eye (pink), yolk (yellow), tail fin (light blue) and fluorescent granules (green). (D) Cartoon showing the location of HSPCs in the CHT in the tail at 3dpf. (E) Correlation of HSPC counts in Tg(itga2b:GFP) embryos at 3dpf between manual counts and granule counts using the *Athena* Zebrafish application, analysed using simple linear regression with r = 0.844.

IDEA Bio-Medical developed a novel Zebrafish image analysis application for their WiSoft^®^ *Athena* software package that identifies the fish and internal anatomy with no required user input. This analysis application uses a unique deep-learning AI algorithm to analyse the brightfield images of full zebrafish (see Methods), trained utilizing hundreds of zebrafish images as input, in conjunction with multiplexed fluorescence channel analysis. The software was trained to use brightfield images to automatically detect the outline of zebrafish embryos, while concomitantly identify internal anatomical structures and regions (Fig. 1B). Currently, the internal anatomy detected includes: eye, heart, tail fin, yolk sac, spine, bladder and otic vesicle (Fig. 1B). The embryo body is also subdivided into the head, trunk and tail regions. Each segmented region can be analysed for features such as morphology (*i.e.* eye size) or count (*i.e.* count 1 vs. 2 eyes to determine orientation), and is combined with fluorescence quantification including intensity metrics and identification of labelled structures, such as fluorescent cells or granules (Fig. 1C).

This analysis approach also permits the automatic selection of wells containing, exclusively, the desired side-oriented fish (illustrated in Fig. 1D), without manual image inspection. Although the use of the alignment plate improves the number of fish in the correct orientation for analysis (Fig. 1A,B), a small number of fish in each plate are not properly aligned in the well (lying on their back, tail out of viewing window etc., (Fig. S2)). This is usually <5% per plate. The side-orientated fish were selected using anatomical attributes identified by the *Athena* software, specifically those having both an eye count and a tail count equal to one, thereby excluding empty wells and fish in undesirable orientations. Defining a minimum fish and tail area in the software also excludes wells where the fish is only partially in the well (see Methods). Automatic exclusion of these wells without any manual inspection of the images is a critical feature for high throughput applications and avoids skewing statistics through incorrect or biased sample selection. While these improperly oriented fish could be correctly oriented via manual intervention, the required hands-on time far exceeds that needed to prepare, scan, process and analyse an additional 96 well plate and results in a lower total amount of useful data.

The rapid image acquisition combined with batch image processing and automatic detection of the fish and its anatomy allows for high throughput analysis with minimal user input. The variety of features detected and the rapid, easy-to-use customisability of the application makes this platform a highly versatile tool for a number of screening applications, as we detail below.

### Accurate counts HSPCs in the tail of 2-4dpf zebrafish embryos

The primary goal of our screen is to accurately enumerate HSPCs, using the Tg(itga2b:GFP) reporter line (Fig. 1C). At 3dpf the HSPCs reside in the caudal haematopoietic tissue (CHT, analogous to fetal liver) in the base of the tail and in close proximity to the caudal-most tip of the yolk sac (Fig. 1D). We aimed to develop a screening platform that would provide an automatic count of the number of cells exclusively in this region to allow us to analyse the effects of myelodysplastic syndrome (MDS)-related genetic mutations that have been identified in the HSPCs of patients. Ultimately such a tool will enable a large-scale small-molecule library screen to target mutant stem cells, to identify drugs that exclusively deplete the mutant HSPC population. One of the challenges in automating our assay is that zebrafish at 3dpf have substantial auto-fluorescence, particularly around the head and yolk (Fig. 1C). The HSPCs of interest are the GFP^lo^ cells, whilst the GFP^hi^ cells are thrombocytes, some of which are in circulation (Lin et al., 2005). This meant that simply thresholding out the auto-fluorescence in the whole image erroneously eliminated a large number of cells of interest, or the inclusion of fluorescent spots not relevant to our question. The *Athena* zebrafish application can provide an automatic count of the fluorescent granules in a chosen region of the fish, in our case the tail (Fig. 1C, outlined in red), without manual inspection or segmentation of the images. Analysis of a 96-well plate takes around 10 minutes. To verify the counts measured, we analysed a 96-well plate of wild-type transgenic zebrafish at 3dpf with the *Athena* software and compared results to a manual cell count in the same images. The two counting methods were strongly correlated, r=0.844 (Fig. 1E), indicating that automated detection provides equivalent results to manual counting.

Stem cells first migrate to the CHT around 2dpf where they then start to proliferate (Chen and Zon, 2009). We tested our platform on embryos of different ages to see the accumulation of HSPCs in the CHT over time (Fig. 2A). Using our automated HSPC counting assay, we confirmed that small numbers of HSPC are present in the CHT at 2 dpf and this steadily increases over the next 48 hours (Fig. 2A-B). We also attempted to image embryos at 5dpf but due to the inflation of the swim bladder, very few fish aligned in the desired side-orientation and our analysis could no longer be carried out with the desired throughput.

**Figure 2.**
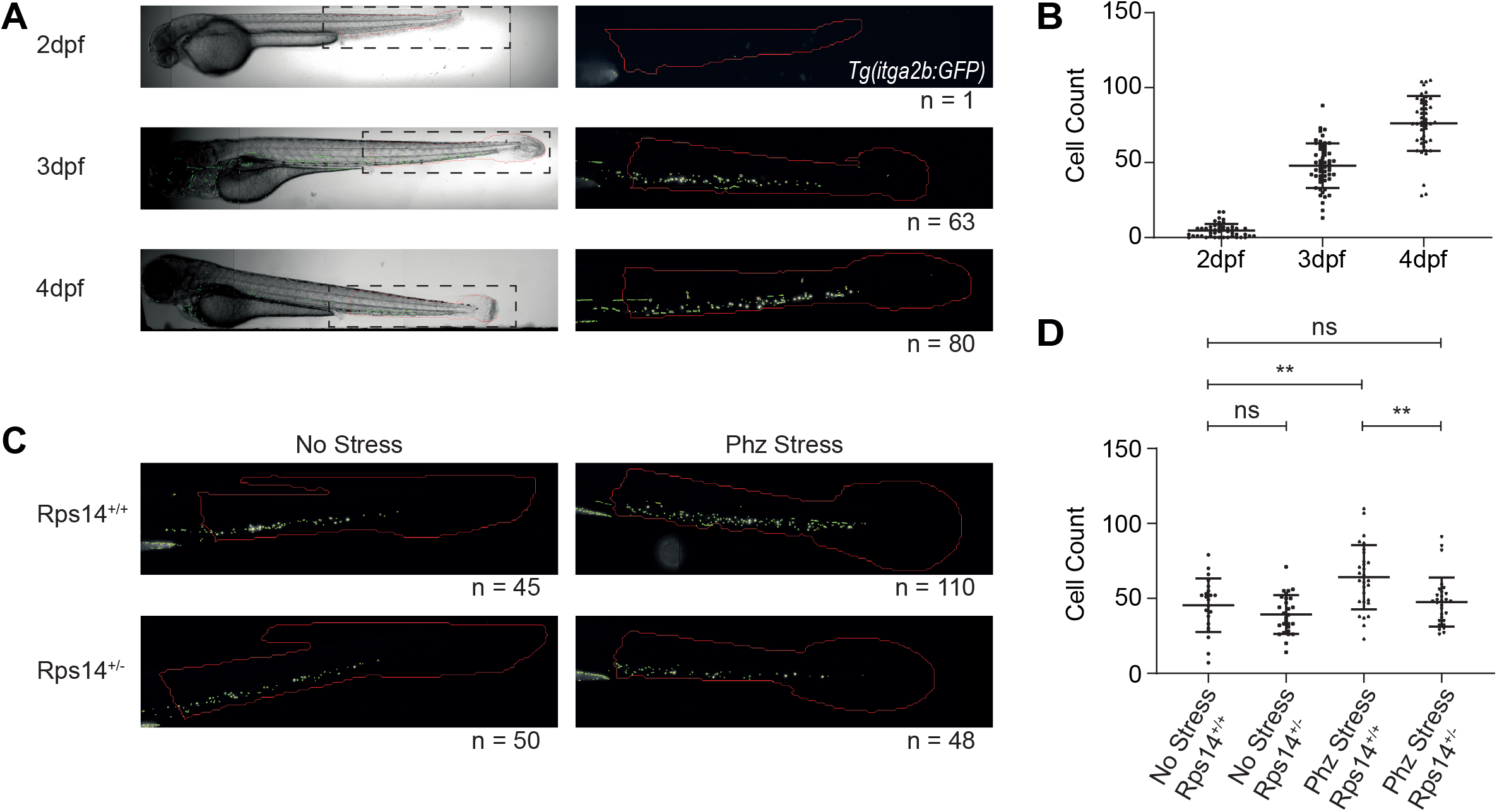
(A) Brightfield and fluorescent images of Tg(itga2b:GFP) embryos at 2-4dpf and analysed for HSPC count (green) in the tail region (red). (B) HSPC counts at 2-4dpf showing the increase in HSPC number over time. (C) Fluorescent images of the tail of Rps14^+/+^ and Rps14^+/−^ Tg(itga2b:GFP) at 3dpf following phz haemolytic stress for 24h at 24hpf alongside unstressed controls analysed for HSPC count (green) in the tail region (red). (D) HSPC count of the different conditions showing an increase in HSPC count in the phz stressed Rps14^+/+^ embryos, which does not occur in the Rps14^+/−^ embryos (Peña et al., 2020). Statistical analysis using unpaired t-tests.

### Detection of phenotypic differences in HSPC and myeloid cell count

We have developed several models for MDS in Zebrafish, including an Rps14 mutant line (Peña et al., 2020). In this model, heterozygous embryos are phenotypically indistinct from wildtype animals unless the animals are subjected to stress. Phenylhydrazine (phz) is used as a haemolytic stress in zebrafish, causing oxidation of haemoglobin and anaemia (Lenard et al., 2016, Shafizadeh et al., 2004, Ferri-Lagneau et al., 2012). If phz is applied for 24 hours at 24-48hpf, only the wildtype embryos recover from the induced anaemia (Schneider et al., 2016, Peña et al., 2020). Using our HSPC counting assay, we showed that at 3dpf, phz leads to an increase in HSPCs at 3dpf in the wild types in response to anaemia, a response not observed in the Rps14 mutants (p < 0.01, Fig. 2C-D (Peña et al., 2020)).

To further define the sensitivity of the platform we assessed the effects a range of non-lethal doses of X-ray irradiation on HSPC numbers in the embryo (Traver et al., 2004, McAleer et al., 2005). Embryos were irradiated at 2dpf before imaging at 3dpf. This led to a significant reduction in stem cells with 40 Gy and a further reduction with 100 Gy (Fig. 3A). The platform allows for swift analysis of these images in large numbers, and the phenotypic differences are easily measured with high statistical significance (p < 0.001, Fig. 3B).

**Figure 3.**
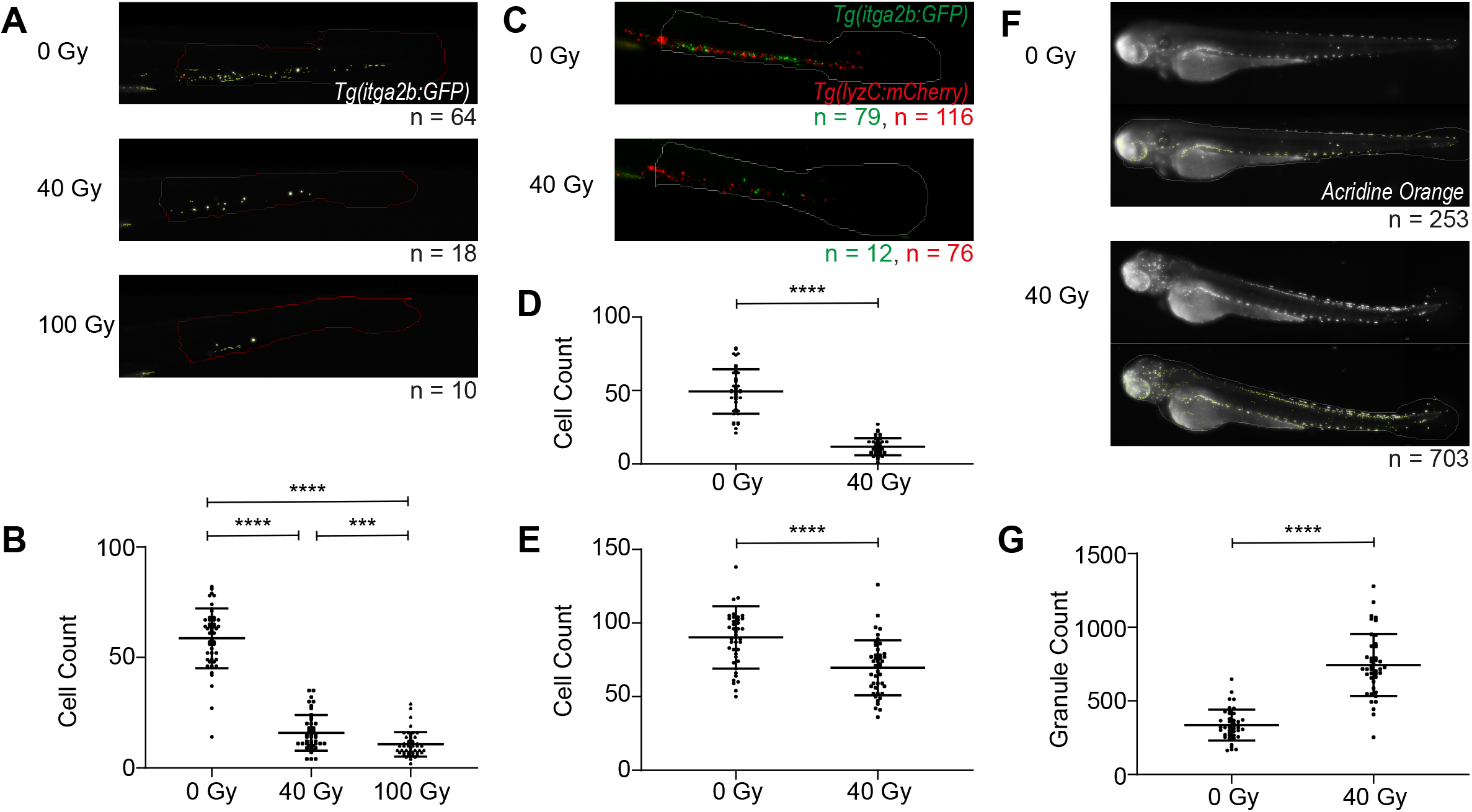
(A) Fluorescent images of the tail of Tg(itga2b:GFP) embryos at 3dpf, irradiated at 2dpf with 0 Gy, 40 Gy or 100 Gy x-ray analysed for HSPC count (green) in the tail region (red). (B) HSPC counts showing the reduction in HSPC number with increased irradiation. (C) Fluorescent images of the tail of Tg(itga2b:GFP)(LyzC:mcherry) dual fluorophore embryos at 3dpf irradiated at 2dpf with 0 Gy or 40 Gy x-ray analysed for HSPC (green) and myeloid (red) in the tail (white). (D) HSPC count of GFP-positive cells and (E) myeloid cell count of mCherry-positive cells in the tail showing the decrease of both cell types with irradiation. (F) Fluorescent images of acridine orange stained embryos at 3dpf following irradiation at 2dpf with 0 Gy or 40 Gy x-ray analysed for granule count (green) in the whole fish. (G) Granule counts in the whole fish showing an increase in apoptosis following irradiation. Statistical analysis using unpaired t-tests.

Beyond analysing solely HSPCs in our MDS models, we aimed to develop a versatile screening platform that we could easily extend to other assays including other zebrafish transgenics to label different cell types. We performed multiplexed fluorescence image acquisition using a double colour transgenic line with Tg(itga2b:GFP) and Tg(lyzC:mCherry), which has mCherry-tagged myeloid cells (Buchan et al., 2019), for imaging in both the red and green channels (Fig. 3C). We used this fish line in the X-ray irradiation assay and identified both the GFP-tagged HSPCs and mCherry-tagged myeloid cells in the same fish (Fig. 3C) by imaging in both fluorescent channels and applying the fluorescent spot counting functionality to each of the two colours individually. Concomitant with the decrease in GFP-tagged HSPCs we also observed a decrease in mCherry-tagged myeloid cells after irradiation (Aldridge and Radford, 1998, Lehnert et al., 1985) (p < 0.0001, Fig. 3D-E).

### Extended applications using fluorescent output – apoptosis and hair cell detection

We wished to further extend the utility of our platform beyond haematological-related screens, to other assays that are commonly used for drug screening in zebrafish embryos, showing the broad applicability to a variety of screening applications. Apoptosis assays in zebrafish can be utilised for drug screens, including analysis of regulated and induced apoptosis and for assessing toxicity (McGrath and Seng, 2013, Parng et al., 2004). Acridine orange is a fluorescent apoptosis marker that can easily be used to visualise cell death in zebrafish by incubation of live embryos in the stain followed by fluorescence imaging (Abrams et al., 1993, Furutani-Seiki et al., 1996, Tucker and Lardelli, 2007). Rapid automated quantification of this fluorescence could be a valuable tool for drug screening and toxicity assessment of chemical screens. To test if our screening platform could be adapted to assess cell death, X-ray irradiation at 2dpf was used to induce apoptosis in zebrafish embryos before analysis at 3dpf. Irradiation in zebrafish embryos leads to apoptosis, with particularly sensitivity towards cells within the spinal cord (Geiger et al., 2006). We labelled dying cells post irradiation with the supravital stain acridine orange and imaged on the *Hermes* in high-throughput (Fig. 3F). The *Athena* analysis pipeline was optimised by adjusting the threshold, smoothing and area parameters of the granule detection, to count the small stained granules which define the dying cell population in the whole fish. The increased cell death with irradiation was easily observed (p < 0.0001, Fig 3G).

Assays analysing hair cells in the lateral line in zebrafish have also been utilised in drug discovery for compounds that mediate deafness and ototoxicity due to the similarities between these cells and the inner ear hair cells in humans (Whitfield, 2002, Nicolson, 2005, Ton and Parng, 2005). Hair cells in the lateral line are arranged into neuromasts along the head and body of the zebrafish and can be stained using fluorescent marker YO-PRO-1. This has been utilised in several chemical and genetic screens (Chiu et al., 2008, Vlasits et al., 2012, Esterberg et al., 2013, Pei et al., 2018, Owens et al., 2008). To date these screens have required manual evaluation of the neuromasts either by counting or scoring for degradation. We therefore sought to replicate the results of one of these screens using our automated analysis platform. Chiu et al. screened 1040 FDA approved compounds for ototoxicity, and found pentamidine isethionate (PI) and propantheline bromide (PB) reduced hair cell survival by almost 50% at 100 μM (Chiu et al., 2008). We incubated embryos at 4dpf with 100 μM PI or PB for 1 hour before staining with YO-PRO-1 and imaging on the *Hermes*. Both compounds led to a reduction in hair cells (Fig. 4A). *Athena* analysis of fluorescent granules contained within the whole fish were used to define regions of interest (the neuromasts) and measure their fluorescence intensity and area to examine the degradation of hair cells therein. Both compounds led to a statistically significant reduction in both integrated fluorescence intensity and fluorescent granule size (Fig. 4B-C, p < 0.0001), in agreement with the results from Chiu et al.

**Figure 4.**
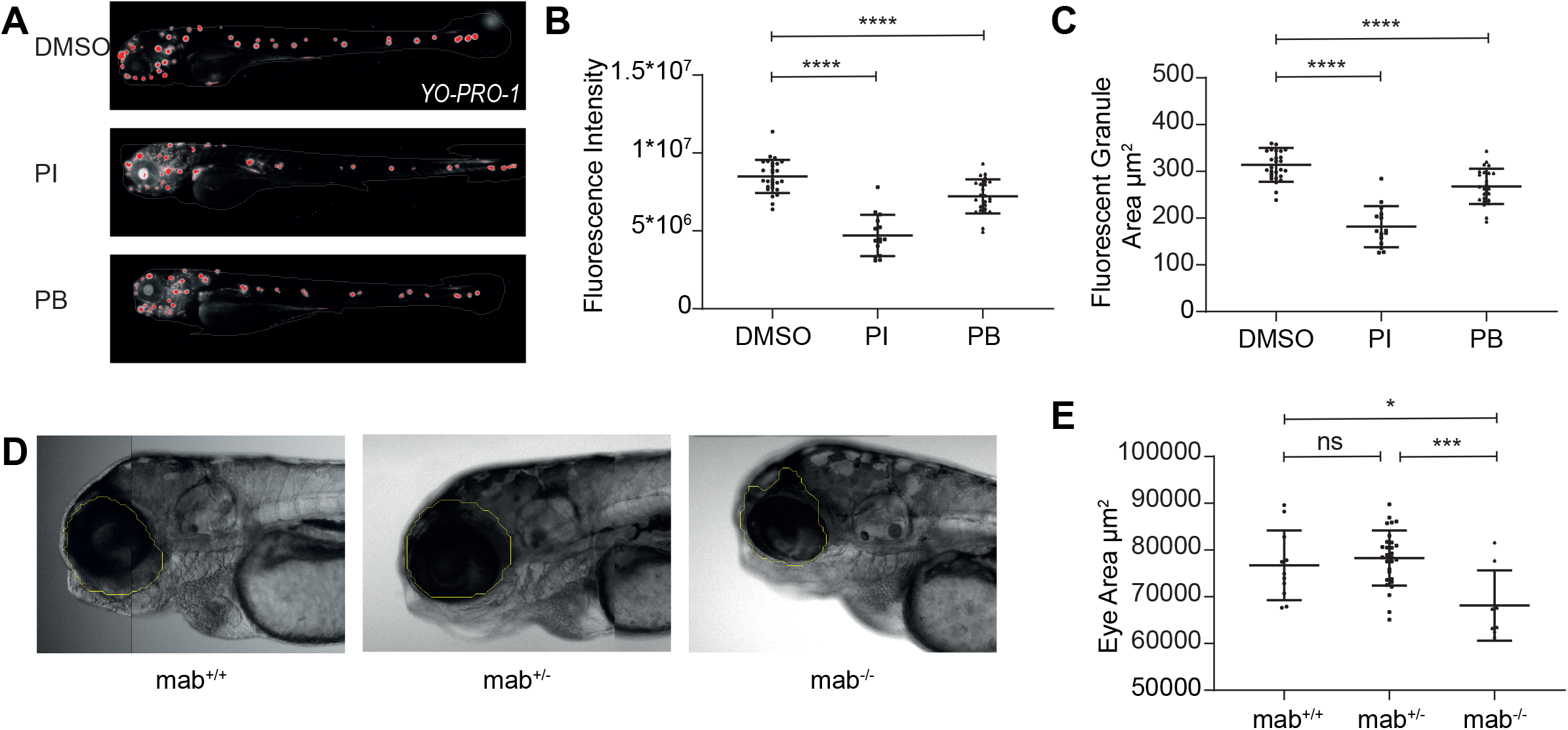
(A) Fluorescent images of YO-PRO-1 stained embryos at 4dpf following treatment for 1 hour with DMSO, 100μM pentamidine isethionate (PI) or 100μM propantheline bromide (PB) analysed for fluorescent granules (red) in the whole fish. (B) Total fluorescence intensity in the whole fish and (C) average area of the fluorescent granules, showing decrease with drug treatment as previously described (Chiu et al., 2008). (D) Brightfield images of mab21l2^+/+^, mab21l2^+/−^ and mab21l2^−/−^ mutant embryos at 4dpf analysed for eye size (yellow). (E) Eye size showing the reduced eye size in mab21l2^−/−^ mutants. Statistical analysis using unpaired t-tests.

### Brightfield phenotypic analysis – eye size

In addition to fluorescence assays, we assessed if our platform could be adapted for morphological screens in brightfield. Mab21l2 is involved in healthy eye development, and zebrafish carrying homozygous *mab21l2* mutations display microphthalmia (smaller eyes) (Gath and Gross, 2019, Deml et al., 2015, Hartsock et al., 2014). Using an incross of *mab21l2*^*u517*^ heterozygous embryos (Wycliffe et al., 2020), imaged in brightfield with the *Hermes* at 4dpf, we used *Athena* to identify they eye in brightfield and measure the size. Consistent with previous studies, *mab21l2*^*u517*^ homozygous are smaller in size (Fig. 5D), as measured using *Athena* (p < 0.05, Fig. 5E).

## Discussion

Zebrafish embryos provide an excellent opportunity for *in vivo* screens, but often such screens rely on time-consuming manual assays or the development of bespoke automated solutions that require time and expertise to develop. Utilising the *Hermes* and *Athena* HCS platform from IDEA Bio-Medical offers an automated solution for zebrafish screens, which is both versatile and easy to use.

Image acquisition is straight-forward to set-up, with simple plate loading and imaging. The plates are loaded in the lab with a standard pipette with no special treatment of the embryos required beyond the use of the alignment plate. Imaging is easy to carry out – the plate is placed in the microscope and the parameters chosen using user-friendly software, where changes can be visualised interactively before imaging. Imaging parameters can be saved and loaded to provide consistency between images acquired at any time. Unlike the VAST system which offers 360° imaging of embryos in a capillary (Pulak, 2016, Pardo-Martin et al., 2010, Pardo-Martin et al., 2013, Chang et al., 2012), the *Hermes* microscope images the fish from beneath the plate. This allows for side-on multiplexed images of fish at 2-4dpf (Fig. 2A) in multiple fluorescence colours (Fig. 3C). For many applications this is sufficient, and although not all embryos are perfectly aligned in the wells with our method, this is limited to around 5% per plate and the software parameters permit automatic exclusion of these wells without manual inspection of the images.

The *Athena* Zebrafish application allows for easy, automated analysis of the acquired images. We have demonstrated one of the principle advantages of this system is the ease with which customisation to different assays can be undertaken. Following set up with the HSPC assay, each of the subsequent optimisations shown in this manuscript were completed in a single experiment. We have shown that the system can accurately replicate published results from a number of different assays: HSPC counts in stressed and unstressed rps14 mutant embryos (Peña et al., 2020) (Fig. 2C-D), hair cell degradation in drug-treated embryos (Chiu et al., 2008) (Fig. 4A-C) and reduced eye-size in mab21l2 mutants (Wycliffe et al., 2020) (Fig. 4D-E). Acquisition and analysis protocols can be customised and then saved, allowing for consistency between experiments, an asset for high-throughput screens carried out over weeks or months.

Both image acquisition and analysis are extremely fast once the protocols have been saved. For a 96-well plate of embryos, image acquisition takes around 15 minutes, image processing a further 20 minutes and image analysis around 10 minutes. The ability to screen and analyse large numbers of embryos in a single day with high consistency and such limited user input makes this platform particularly useful for high-throughput applications such as drug and genetic screens, where large numbers of embryos are required.

Zebrafish provide a unique opportunity for drug screening in whole live animals since they produce large numbers of fast developing, transparent embryos. The development of an automated screening platform that is fast, customisable and easy to use without specialist knowledge or training will allow for much better utilisation of this unique system. As shown in this manuscript, the platform can be applied to a large number of different assays across a large range of research topics. The speed of acquisition and analysis allows access to larger compound libraries, the consistency between experiments allows for comparison of results across multiple experiments, and the ease of use makes the platform accessible to all. The training set used to identify fish and anatomy are centralized and continuously updated/maintained which also provides reproducibility and consistency between research groups.

The utility of this platform extends beyond its intended role as a screening platform to high-content quantitative analysis across other more targeted experiments massively increasing experimental throughput and statistical confidence. The ease of use, speed of imaging and analysis, and consistency between images acquired at different times makes this platform useful for both small and large-scale experiments. We now plan to use this platform for a small-molecule drug screening, using the automated HSPC count presented here to conduct a synthetic lethal screen in an MDS-related mutant background.

## Materials and Methods

### Zebrafish Husbandry and Experimental Conditions

Zebrafish (Danio rerio) stocks were maintained according to standard procedures in UK Home Office approved aquaria (Westerfield, 2007). Embryos were obtained from wild-type AB or AB/TL, transgenic strains Tg(itga2b:GFP) which has green-fluorescent HSPCs (Lin et al., 2005) and Tg(Lyzc:mcherry) which has red-fluorescent myeloid cells (Buchan et al., 2019) or mutant strains *rps14*^*E8fs*^ (Peña et al., 2020) and *mab21l2*^*u517*^ (Wycliffe et al., 2020). Embryos were staged according to Kimmel et al. (Kimmel Charles et al., 1995) and expressed in hours/days post-fertilization (hpf/dpf). All procedures complied with Home Office guidelines.

### Preparation of Embryos for Imaging

At 24hpf, embryos were dechorionated using pronase, and raised in 0.003% phenylthiourea-supplemented E3 medium to prevent pigment formation. At 2-4 dpf embryos were anaesthetised with tricane, and then loaded into a 96-well zebrafish alignment plate (Hashimoto ZF plate, Japan, 96-well) in 75 μL of the E3 medium using a wide orifice tip. The plates were then gently centrifuged at 200 rcf for 20 seconds before imaging.

### WiScan^®^ *Hermes* Image Acquisition

The 96-well plates were imaged using the WiScan^®^ *Hermes* High Content Imaging System (IDEA Bio-Medical, Rehovot, Israel). Images were taken in brightfield and green and red fluorescence channels at a 4x magnification with a well coverage of 150% and field density of 150%. This gave 4 overlapping images along each well. Brightfield was imaged with 35% light intensity, 40 ms exposure and 30% gain. Fluorescence channels were imaged with 90% light intensity, 200 ms exposure and 30% gain. Z-stacks were taken in 5 planes with an inter-plane distance of 50.6 μm.

An accompanying image pre-processing software package (Advanced Data Processing software, IDEA Bio-Medical, Rehovot, Israel) was used to process the raw images of multiple data sets in batch prior to quantitative analysis. This software loads one or more image data sets obtained on the WiScan^®^ *Hermes* microscope, then performs one or more sequential image processing operations in batch. Image processing operations can include intensity projection (through time or Z-slices), fluorescence deconvolution, sharpest (most in-focus) Z-plane selection, and image montage (stitching). Here, the sharpest z-slice for each raw field of view in the brightfield channel was selected, and maximum intensity projection for each raw field of view in the fluorescence channel was performed. Subsequently, all fields of view (sharpest brightfield and maximum fluorescence) in a well were stitched together to provide a single, 2-colour channel image per well.

### WiSoft^®^ *Athena* Image Analysis

Image quantitation was performed with the WiSoft^®^ *Athena* software *Zebrafish Application* (IDEA Bio-Medical, Rehovot, Israel). The software application performs automated, multiplexed image analysis by processing brightfield and fluorescence channels simultaneously, with different algorithms. Brightfield analysis of the whole zebrafish embryo utilized a novel deep learning based artificial intelligence algorithm to identify the fish and internal structures. The method is based on a supervised convolutional neural network and was trained using hundreds of images of 2-4 dpf zebrafish embryos as input, each paired with manually segmented ground-truth outputs. An option for custom, manual segmentation is also present in the *Zebrafish* application. The resulting algorithm automatically identifies the fish outline, internal anatomical structures and three fish regions (head, trunk, and tail). Users input only minimum and maximum size constraints to select the appropriate range for anatomical objects/regions that are to be identified and analysed. Fluorescence channels are analysed using image analysis techniques with optimized algorithms (smoothing, background subtraction, thresholding and area constraints, Table 2) to identify high signal-to-background objects, such as fluorescent granules.

The morphological features (length, area, shape, etc.) of the fish, organs and regions are quantified based on the structures identified in the brightfield channel. Objects (*i.e.* fluorescent granules) identified in the fluorescence channels that are contained within the fish outline are included in quantification, while spots outside the fish are disregarded. Fluorescent spots are quantified regarding their count and intensity within the fish as a whole, as well as within anatomical or regional structures (e.g. granule count within the tail, spot intensity within the spine, etc.).

Parameters were set to define the permitted size of detected fish and internal features in 2-4dpf embryos (Table 1). No additional parameter input was required to identify these objects, since the AI algorithm was trained to identify them automatically. A population of on-side orientated fish for analysis was defined as those with an eye count and a tail count equal to one.

**Table 1.** *Athena* Zebrafish Application parameters for detecting an embryo within the well.

| Feature Detected | Minimum Area ($\mu\text{m}^2$) | Maximum Area ($\mu\text{m}^2$) |
| --- | --- | --- |
| Fish | **950000** | 1600000 |
| Head | **15000** | 1000000 |
| Trunk | **25000** | 1300000 |
| Tail | **150000** | 1000000 |
| Eye | **20000** | 250000 |
| Yolk | **100000** | **300000** |
| Fin | **15000** | 300000 |

**Bold** text shows parameters that have been adjusted from the default

For fluorescence analysis, granules were identified within the *Athena* Zebrafish Application by defining the fluorescence threshold, area of the granules, background subtraction and smoothing. Parameters were customised for each assay (Table 2).

**Table 2.**
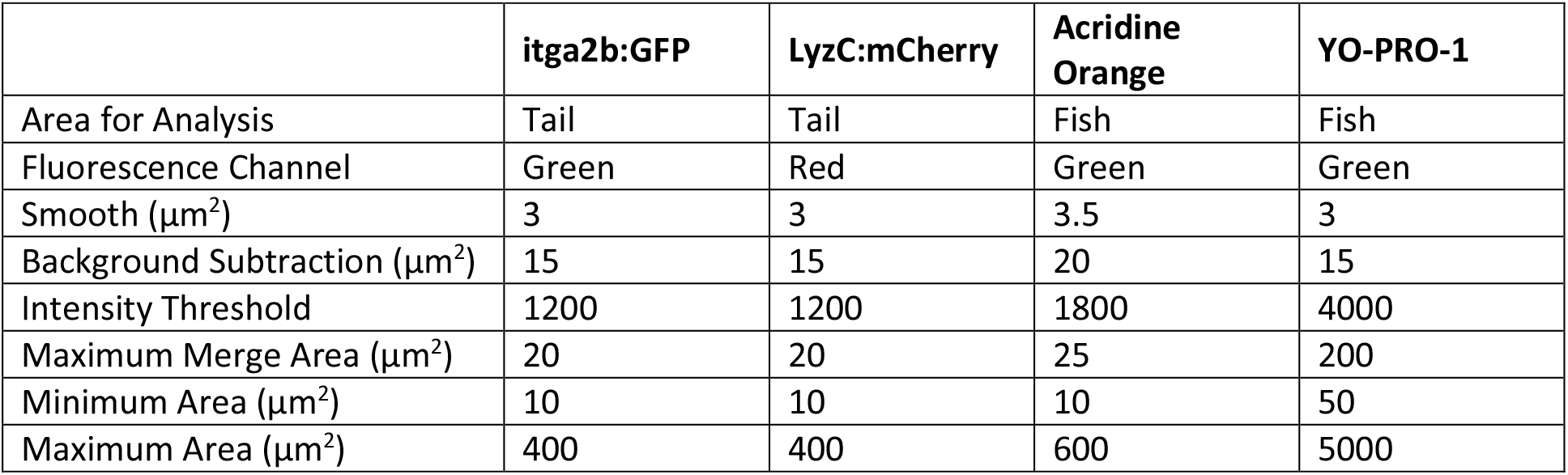
*Athena* Zebrafish Application parameters for fluorescence granules for different assays.

### Genotyping Mutants

For Rps14 and mab21l mutants, genotyping was required post imaging. Embryos were removed in 50 μL of nuclease-free water using a multi-channel pipette and wide orifice tips and added directly to 1 μL 50x HotSHOT base solution (KOH 1.25 M, EDTA 10 mM). After incubation at 95°C for 30 minutes the solution was neutralised with 1x HotSHOT neutralising solution (40 mM Tris-HCl). To identify the genotype of Rps14 and mab21l mutants we used the Kompetitive Allele Specific PCR genotyping assay (KASP, LGC Genomics).

### X-Ray Irradiation

At 48hpf embryos were transferred to 6-well plates for irradiation. X-rays (250kV, 12.5mA, 1.0 mm Al filter) for a total dose of at 40 Gy or 100 Gy using an AGO HS 320/250 X-ray machine (AGO X-ray Ltd.) equipped with a NDI-321 stationary anode X-ray tube (Varian). Embryos were placed back at 28°C for 24 hours before imaging at 3dpf as above.

### Acridine Orange Staining for Detection of Apoptosis

At 3dpf, live embryos in 6-well plates were incubated in acridine orange (Invitrogen) staining solution (1 μg acridine orange in E_3_ medium) in the dark for 30 minutes with gentle rocking. Embryos were swiftly washed with E_3_ medium four times before being anaesthetised with tricaine, and then loaded into plates for imaging as above. Plates were kept in foil to shield from light and imaged promptly after staining.

### Hair Cell Assay

Protocol adapted from Chiu et al. (Chiu et al., 2008). At 4dpf, live embryos in 6-well plates were incubated in YO-PRO-1 (Invitrogen) staining solution (2 μM YO-PRO-1 in E_3_ medium) in the dark for 30 minutes with gentle rocking. Embryos were swiftly washed with E_3_ medium four times. Embryos were then treated with 100μM pentamidine isethionate (PI), 100μM propantheline bromide (PB) or DMSO control for 1 hour. Embryos were then anaesthetised with tricaine and loaded into plates for imaging as above.

---

**Figure S1.**
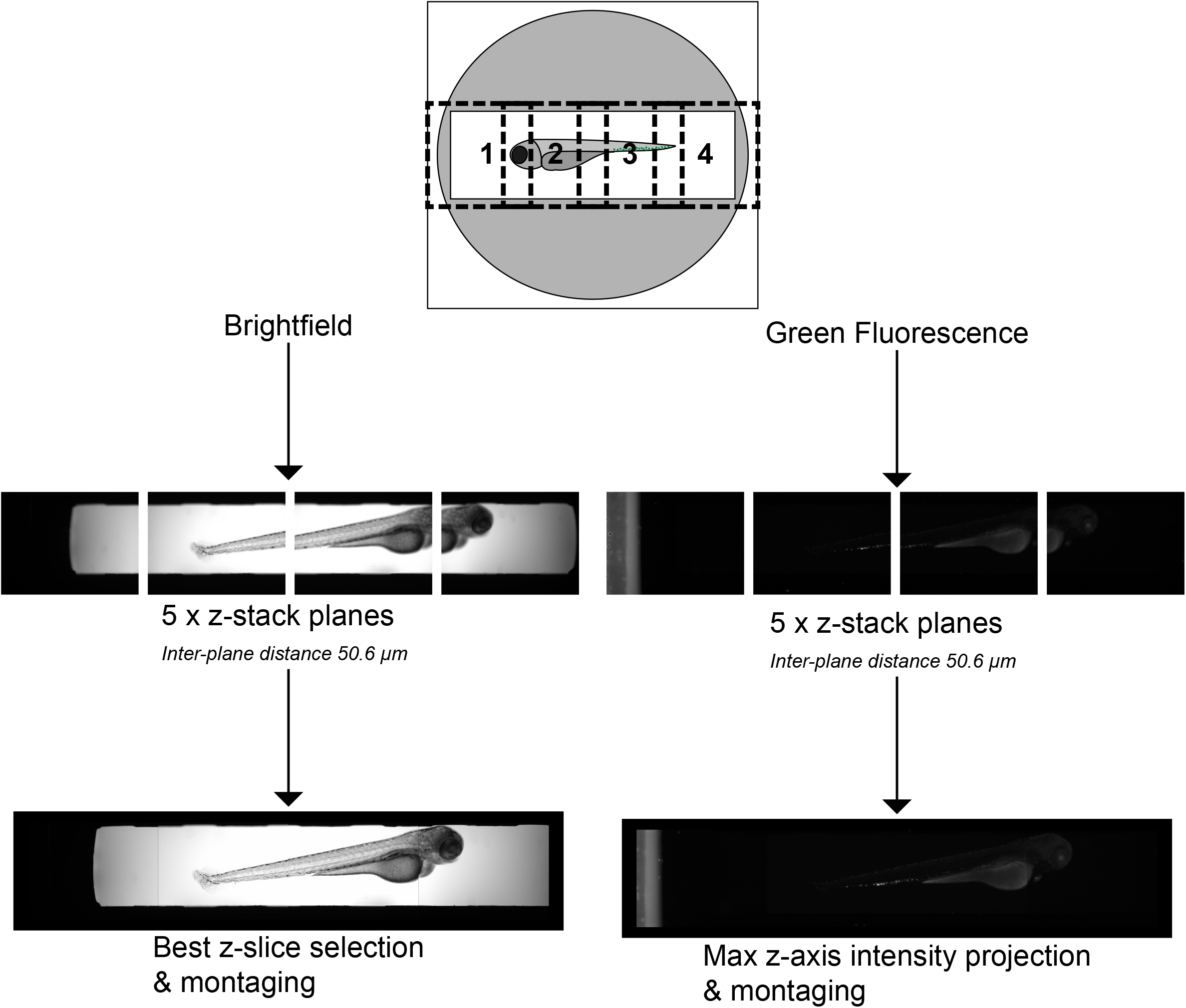
Illustration of image pre-processing from raw images. Embryos were imaged using a 4X objective and Z-stack acquisition of five slices, with four overlapping images along each well in brightfield and fluorescent channels. An accompanying image pre-processing software package (Advanced Data Processing software) was used to process the raw images selecting the sharpest z-slice for each raw field of view in the brightfield channel was selected, and maximum intensity projection for each raw field of view in the fluorescence channel. All fields of view (sharpest brightfield and maximum fluorescence) in a well were then stitched together to provide a single image in each channel per well.

**Figure S2.**
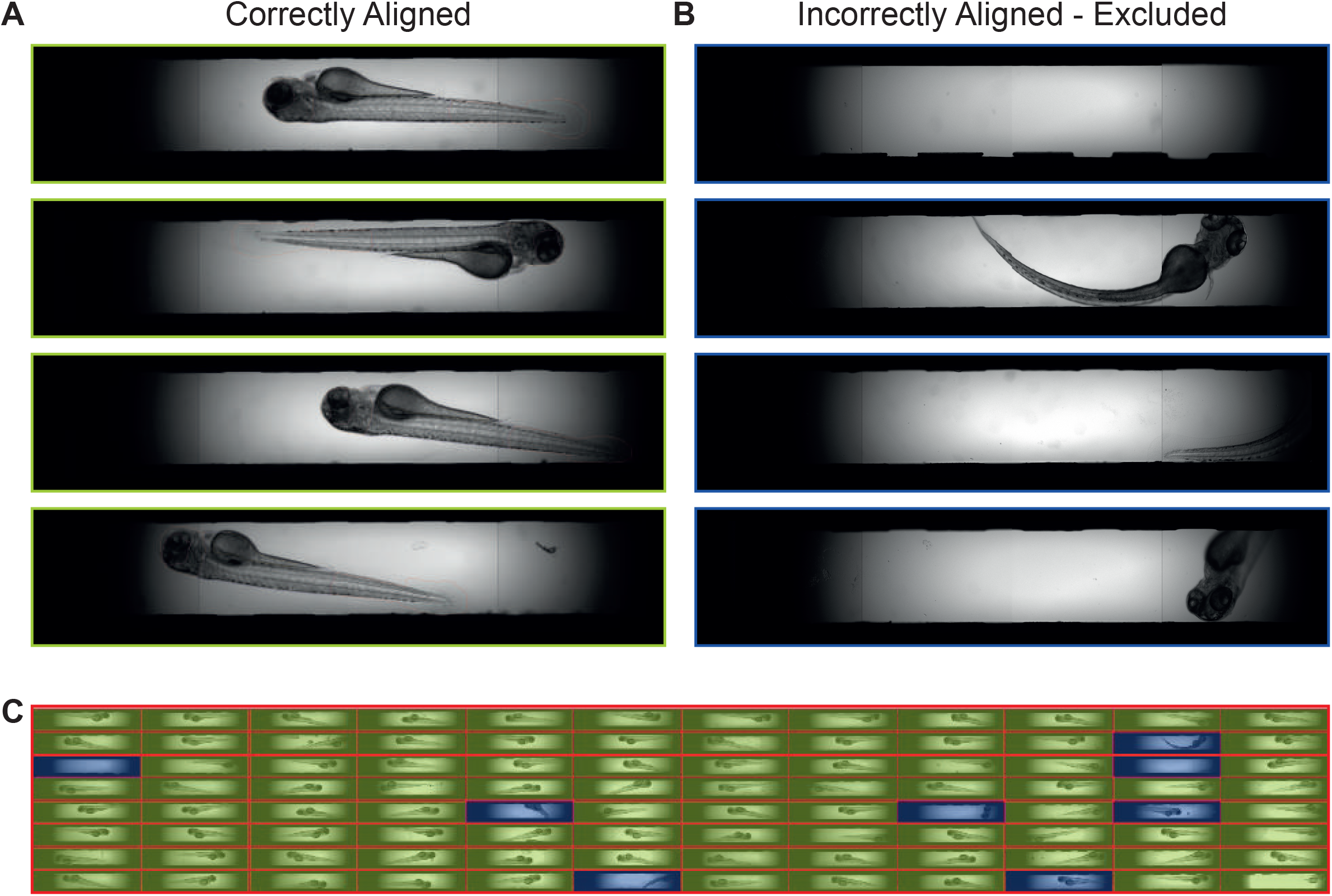
Wells containing fish not properly aligned in the well (lying on their back, tail out of viewing window etc.) rather than in the desired orientation are automatically excluded by the *Athena* software. This is usually <5% per plate. The side-orientated fish were selected using anatomical attributes identified, specifically those having both an eye count and a tail count equal to one, thereby excluding empty wells and fish in undesirable orientations.

